# Gene regulatory effects of a large chromosomal inversion in highland maize

**DOI:** 10.1101/861583

**Authors:** Taylor Crow, James Ta, Saghi Nojoomi, M. Rocío Aguilar-Rangel, Jorge Vladimir Torres Rodríguez, Daniel Gates, Ruben Rellan-Alvarez, Ruairidh Sawers, Daniel Runcie

## Abstract

Chromosomal inversions play an important role in local adaptation. Inversions can capture multiple locally adaptive functional variants in a linked block by repressing recombination. However, this recombination suppression makes it difficult to identify the genetic mechanisms that underlie an inversion’s role in adaption. In this study, we explore how large-scale transcriptomic data can be used to dissect the functional importance of a 13 Mb inversion locus (*Inv4m*) found almost exclusively in highland populations of maize (*Zea mays* ssp. *mays*). *Inv4m* introgressed into highland maize from the wild relative *Zea mays* ssp. *mexicana*, also present in the highlands of Mexico, and is thought to be important for the adaptation of these populations to cultivation in highland environments. First, using a large publicly available association mapping panel, we confirmed that *Inv4m* is associated with locally adaptive agronomic phenotypes, but only in highland fields. Second, we created two families segregating for standard and inverted haplotypess of *Inv4m* in a isogenic B73 background, and measured gene expression variation association with *Inv4m* across 9 tissues in two experimental conditions. With these data, we quantified both the global transcriptomic effects of the highland *Inv4m* haplotype, and the local cis-regulatory variation present within the locus. We found diverse physiological effects of *Inv4m*, and speculate that the genetic basis of its effects on adaptive traits is distributed across many separate functional variants.

**Author Summary:** Chromosomal inversions are an important type of genomic structural variant. However, mapping causal alleles within their boundaries is difficult because inversions suppress recombination between homologous chromosomes. This means that inversions, regardless of their size, are inherited as a unit. We leveraged the high-dimensional phenotype of gene expression as a tool to study the genetics of a large chromosomal inversion found in highland maize populations in Mexico - *Inv4m*. We grew plants carrying multiple versions of *Inv4m* in a common genetic background, and quantified the transcriptional reprogramming induced by alternative alleles at the locus. *Inv4m* has been shown in previous studies to have a large effect on flowering, but we show that the functional variation within *Inv4m* affects many developmental and physiological processes.

**Author Contributions:** T. Crow, R. Rellan-Alvarez, R. Sawers and D. Runcie conceived and designed the experiment. M. Aguilar-Rangel, J. Rodrǵuez, R. Rellan-Alvarez and R. Sawers generated the segregating families. T. Crow, J. Ta, S. Nojoomi, M. Aguilar-Rangel, J. Rodrǵuez D. Gates, D. Runcie performed the experiment. T. Crow, D. Gates, D. Runcie analyzed the data. T. Crow, D. Runcie wrote the original manuscript, and R. Rellan-Alvarez and R. Sawers provided review and editing.

## Introduction

Chromosomal inversions are structural rearrangements that form when a portion of a chromosome breaks in two places and reinserts in the opposite orientation. The reversed order of loci prevent recombination with the non-inverted homologous chromosome, as crossover products are imbalanced and often non-viable [1]. This spontaneous, long-distance genetic linkage is important for speciation and local adaptation because it can capture multiple adaptive and potentially interacting loci in a single haplotype [2–4]. Inversions are common across taxa [1], often pre-date speciation events, and can spread through admixture [5, 6]. They have been linked to adaptive phenotypes and environmental clines [7–9], mating system evolution [10–12], social organization [13], and migratory phenotypes [14].

Chromosomal inversions were first discovered nearly a century ago in *Drosophila* [3, 15] by visualizing karyotypes, and can also be identified based on their effects on recombination rates among nearby markers. However, both techniques are labor intensive and difficult to apply to large-scale population-level surveys within or among species. Modern genome-wide sequencing technologies provide the opportunity to identify inversions more rapidly and comprehensively, leading to the discovery of inversions across a wide range of species [1]. In fact, measured by the number of variable base pairs in a genome, structural variants (e.g. insertions, deletions, duplications, translocations, fusions and inversions) have been shown to account for more variation than SNPs in some species [16].

However, while whole-genome sequencing data can help rapidly discover inversion loci, measure their frequencies across populations, and test for associations with adaptation and speciation, there are still very few examples where the mechanisms that underlie the adaptive value of any particular inversion is known. Are there generally one or two loci of major effect within an adaptive inversion, or do inversions harbor many small-effect variants that combine to make inversion loci highly pleiotropic supergenes [17]? Because inversions suppress recombination across a large genomic region, QTL mapping and Genome-Wide Association Studies have little ability to resolve independent effects of different variants within an inversion locus. This makes fine-mapping nearly impossible. Only in cases of very old inversion loci which have experienced rare recombinants or gene conversion events can association methods or population-genetic signatures of selection successfully identify causal loci within inversions [18]. Alternatively, young inversions may capture adaptive loci, and mapping populations can be created from closely-related, non-inverted individuals [19].

RNA sequencing technologies may be a powerful tool for gaining rapid insight into the evolutionary role of inversion loci, particularly for loci for which the adaptive phenotypes regulated by the locus are unknown. RNA sequencing is a very high-throughput phenotyping technology that can simultaneously measure tens of thousands of different gene expression values from each experimental sample. The expression of each gene responds to a different combination of transcription factors, gene networks and cellular states, so measuring gene expression provides an indirect measurement of a wide range of cellular, developmental and physiological characteristics of an organism. These cellular or physiological traits may be critical for adaptation, yet are often neglected in evolutionary studies because they are difficult and costly to measure directly. At the same time, gene expression analysis can be used to scan across an inversion locus gene-by-gene to identify specific genes that have different cis-regulatory genetic control among alleles. If the functional variation captured by an inversion locus operates by directly altering the expression of genes in the inversion, we can identify these genes by their expression changes without relying on recombination. Together, these two types of gene expression analysis may greatly advance our understanding of inversion loci, particularly those that are not feasible to study by other means.

In this study, we applied population genetic and gene expression analyses to study an inversion locus in maize. Maize is an important crop species worldwide and also a powerful model system for studying the mechanisms of recent and rapid local adaptation. Maize (*Zea mays* ssp. *mays*) was domesticated in the lowland Balsas river valley of southwestern Mexico from the lowland teosinte subspecies, *Zea mays* ssp. *parviglumus*, hereafter *parviglumus* approximately 9000 years ago [20, 21]. Since domestication, populations of maize have been moved into high altitude environments, and landraces collected today show considerable local adaptation to their home elevation in a range of traits [22]. Interestingly, population genetic scans for loci associated with adaptation to elevation gradients have identified several loci common in highland landraces that have been introgressed from a different subspecies of teosinte, *Zea mays* ssp. *mexicana* (hereafter *mexicana*), which occurs in highland environments [23]. One of these introgressed regions, located at approximately 171.7 to 185.9 Mb of chromosome 4, is a chromosomal inversion known as *Inv4m* [24, 25]. *Inv4m* is observed in high altitude locations in Mexico, and is associated with a three day acceleration of flowering time, the largest effect flowering QTL yet found [26]. *Inv4m* also overlaps with a quantitative trait locus (QTL) found in a previous study of leaf pigmentation and macrohairs in teosinte, which are thought to be adaptive in highland environments [27].

We first comprehensively characterized the population-genetic context of the *Inv4m* locus and its association with key agronomic traits using dense whole-genome genotyping data of thousands of Mexican landraces. We found that *Inv4m* is more closely associated with altitude in the center of maize diversity in Mexico than nearly any other locus in the maize genome, and shows clear patterns of antagonistic pleiotropy indicative of a key role in local adaptation. We isolated *Inv4m* from two separate donor sources into a common reference maize background (B73), and used RNA sequencing of samples from nine different tissues and two developmental stages in high and low temperature conditions to identify molecular traits that were differentially regulated by the highland and lowland haplotypess of the inversion. These genes suggest a diverse range of biological processes affected by *Inv4m*, highlighting novel cell-biological or physiological traits that may be involved in maize highland adaptation. Finally, we scanned genes within *Inv4m* for associations with these gene expression traits and identified several outlier genes inside the *Inv4m* locus that are good candidates for further study.

## Results

### *Inv4m* is associated with local adaptation in agronomic traits

To confirm the population genetic signature of local adaptation at *Inv4m*, we determined the *Inv4m* genotype of 4845 maize plants from the SeeD-maize GWAS panel using published unimputed genotype-by-sequencing data. Of these lines, 707 were homozygous for the minor haplotype, and 351 were heterozygous for the locus. Of the 585 plants carrying at least one allele of the minor haplotype and with complete with geographic information, all but 7 were from Mexico, with the majority collected from the central highlands (Fig 1A). Therefore, to assess evidence of local adaptation, we assessed the association of *Inv4m* genotype with elevation among the 1757 Mexican plants. In Mexico, 1186 and 381 plants were homozygous for the alternate haplotype, and 190 were heterozygous, a distribution that significantly differs from Hardy-Weinberg expectations (D=-252.0, p = 1.91e-197). Genotypes at *Inv4m* were strongly associated with elevation, as previously reported [26] Fig 1B. The highland haplotype at *Inv4m* had much lower genetic diversity across the locus; however diversity measures rebounded immediately outside of the published boundaries of the inversion. Plants carrying the highland haplotype did show a slightly lower proportion of segregating sites (*θ*) across chromosome 4 (Fig 1C). Diversity estimates were relatively constant across the “High” haplotype at the *Inv4m* locus, with little evidence of large-scale introgression of lowland alleles; only 5% of markers had segregating variation in common (MAF *>* 0.05) in both haplotypes at the locus, and some of these may have been caused by cryptic paralogous loci which is common in GBS data [28]. The lower diversity estimates are unlikely to be caused entirely by mapping biases against this divergent haplotype. Of the 9201 GBS markers within the locus, *∼* 80% were successfully genotyped in *>* 5% of the highland individuals, and of these markers, 14% were segregating in the highlands and 33% were segregating in the lowlands. Among these markers, rates of missing genotypes were similar between the two alleles. If all of the un-scored markers were actually present and variable in the highland individuals, there would still be fewer segregating positions among highland individuals than lowland individuals.

**Fig 1.**
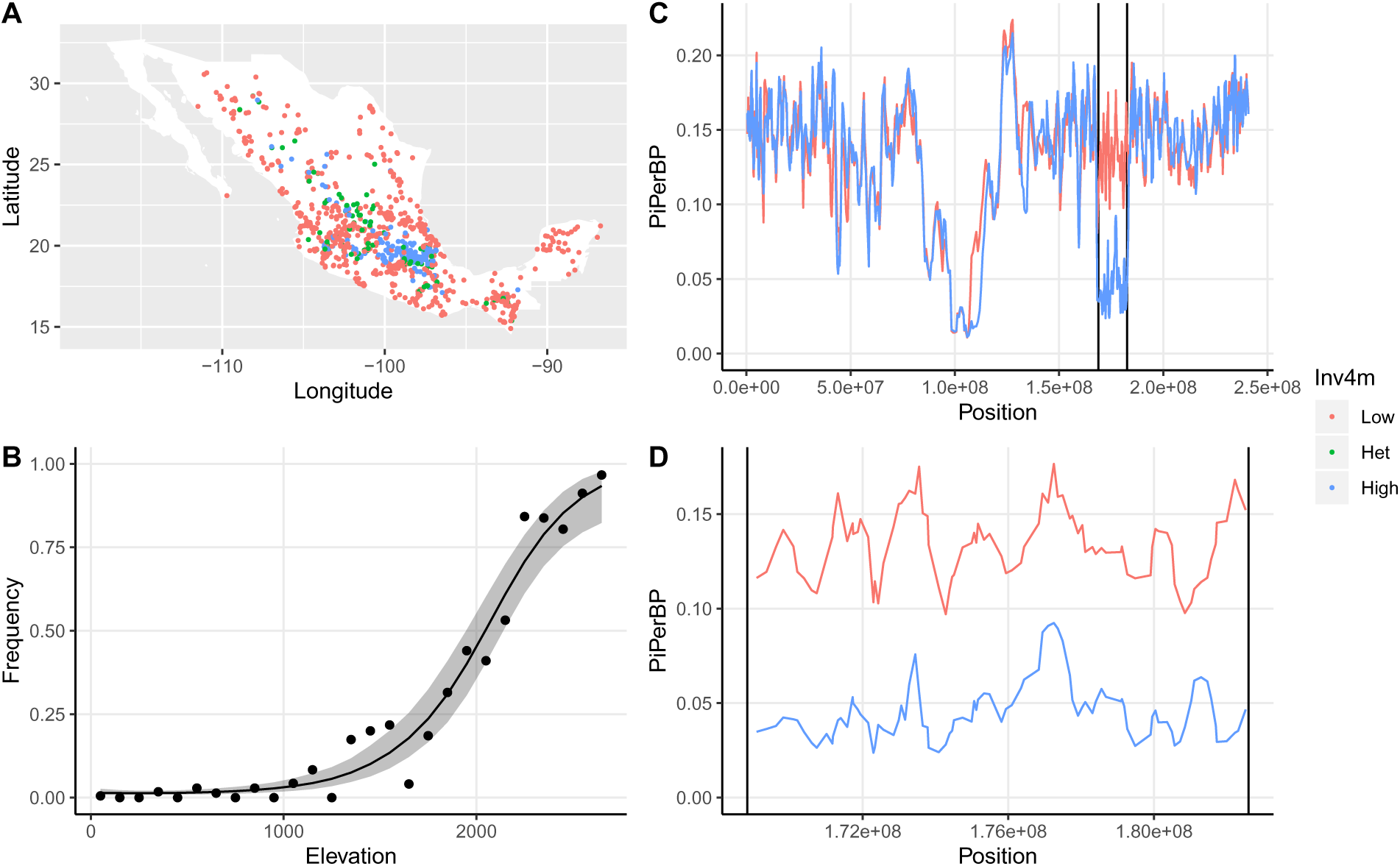
Association of *Inv4m* genotype with environmental factors and agronomic traits. **A**. Geographic locations for each of the 1757 Mexican plants genotyped by GBS, colored by their imputed genotypes at *Inv4m*. **B**. Association of *Inv4m* and elevation. Each point shows the mean frequency of the “High” allele at the *Inv4m* locus among plants from landraces collected in each 100m bin. The ribbon shows a loess fit (±2SE) to the logit-transformed frequencies weighted by the number of landraces in each elevation bin. Bins with fewer than 10 landraces were excluded (those with elevation *>*2700m). **C** and **D**. Diversity estimates *π* and *θ* for 371 sampled plants homozygous for the “Low” haplotype and 371 plants homozygous for the “High” haplotype at *Inv4m*, along chromosome 4 between 162 and 182 Mb. The boundaries of *Inv4m* from [24] are denoted by vertical lines.

Romero-Navarro *et al* [26] identified *Inv4m* as a large-effect QTL in a multi-environment trial of landrace hybrids grown in multiple field sites in Mexico. Gates *et al* [29] analyzed five additional agronomic traits from some of these field trials and found evidence for effects of the *Inv4m* locus on several traits. However, these studies did not explicitly show effect sizes for *Inv4m* across trials or traits, and none of the individual SNP markers in these studies was perfectly associated with our *Inv4m* genotype. Therefore, we re-analyzed the phenotype dataset focusing specifically on estimating the effect of *Inv4m*, and how this effect changed across the elevations of the trials.

The highland haplotype of *Inv4m* was significantly associated with Days-to-Anthesis, the Anthesis-Silking Interval (ASI), Grain Weight-per-hectare, Bare cob weight, and Field Weight, but in each case, the effect size changed across elevations in the direction consistent with local adaptation: earlier flowering, reduced ASI, and increased yield components in highland trials, while the opposite in lowland trials (Fig 2). The highland haplotype was weakly associated with greater plant height, but the relationship was not significant (p=0.11 for the main effect). The relationship between *Inv4m* genotype and these traits was not simply an indirect effect of the change in flowering time; each relationship remained qualitatively the same even after accounting for the the effect of Days-to-Anthesis separately within each trial. However, even though we attempted to account for population structure in our analyses, it is still possible that some of these results remain confounded due to the relatedness among individuals; *Inv4m* is among the markers best-correlated with the elevation of origin, and the first eigenvector of genetic variation across the remainder of the genome (excluding chromosome 4) is also strongly correlated with elevation. Therefore, these results are also consistent with a polygenic basis for the divergence of each of these traits along elevational gradients. Functionally validating the association of traits with *Inv4m* therefore requires experimentally breaking the association between *Inv4m* and the rest of the genome through experimental crosses.

**Fig 2.**
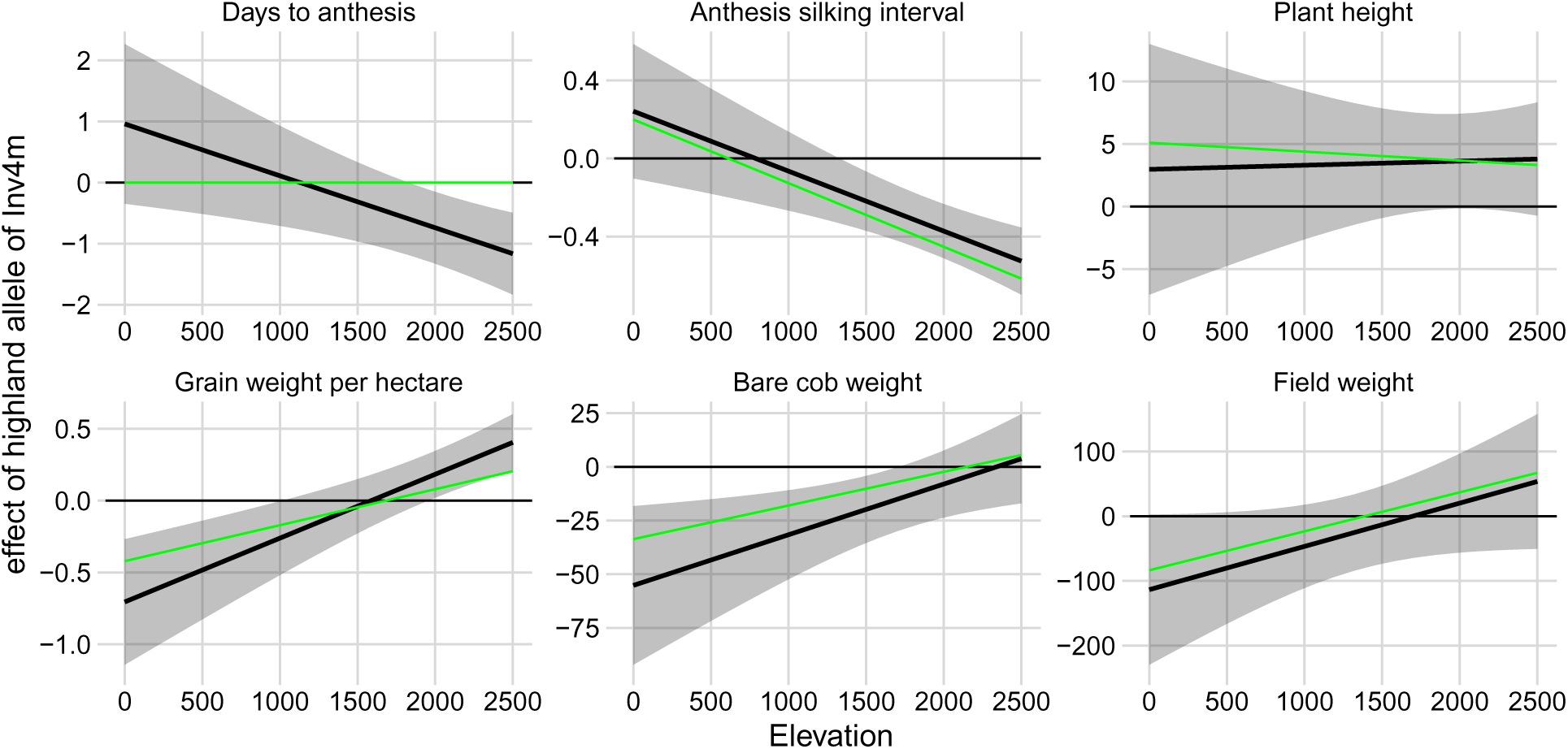
Association of *Inv4m* genotype with agronomic traits depends on trial elevation. We modeled each trait as a function of *Inv4m* genotype, trial elevation, and tester line, with controls for main effects and responses to elevation of the genomic background. Black lines and ribbons show estimates of the effect of the highland allele of *Inv4m* as a function of trial elevation ± 2SE, based on conditional F-tests at the REML solutions of the random effect variance components. Green lines show estimates of the *Inv4m* effect in a model that additionally included effects of Days-to-Anthesis on the focal trait within each trial.

### Isolating *Inv4m* in common genetic background

We generated two sets of segregating families for the *Inv4m* locus by crossing the reference line B73 which carries a lowland haplotype of *Inv4m* to two highland landrace donors (called PT and Mi21). The progeny were backcrossed to B73 for 5 generations and then selfed to generate a population of plants that were largely homozygous B73 across the genome, but segregated for the lowland highland haplotypes at the *Inv4m* locus. We used 81 plants that were homozygous for either the High or Low haplotypes from these families to measure the effect of *Inv4m* on more than 300,000 gene expression traits across 9 tissues (Table 1) and two environments (warm: 32C/22C and cold: 22C/11C).

**Table 1.** Developmental stage and description of tissues sampled for gene expression analysis.

| Developmental Stage | Tissue | Description |
| --- | --- | --- |
| V1 | Root | Primary root |
| V3 | Root | Primary root |
| V3 | SAM | Stem Apical Meristem |
| V1 | Leaf | Pooled leaf tissue |
| V3 | Sheath | Leaf sheath |
| V3 | Leaf base | 5 cm of leaf base |
| V3 | Leaf tip | 5 cm of leaf tip |
| V3 | S2 leaf base | 5 cm of leaf base |
| V3 | S2 leaf tip | 5 cm of leaf tip |

### *Inv4m* has mild effects on seedling emergence

We first tested if *Inv4m* genotype had an effect on seedling emergence time. Emergence times in the cold chamber averaged approximately 9 days, and plants with the High haplotype of *Inv4m* emerged 0.75 and 0.35 days earlier in the PT and Mi21 families, respectively (Table S1).In the warm chamber, the average emergence time was approximately 4 days, and the High *Inv4m* haplotype had no detectable effect (Table S1).

### Quantification of residual landrace alleles in BC_5_S_1_ plants

We used RNAseq reads to identify residual landrace alleles at every expressed gene in each of the BC_5_S_1_ plants. We observed residual alleles in large regions flanking the *Inv4m* locus in plants from all families, despite the 5 generations of backcrossing to B73. In the PT segregating family, residual PT alleles were observed in a 57Mb window surrounding *Inv4m*. In the Mi21 family, residual Mi21 alleles were observed in an 18Mb region surrounding *Inv4m* (Fig 3A). The PT family also segregates for a large paracentric region of residual donor alleles on chromosome 5 and a small region on chromosome 2, and the Mi21 family segregates for a large paracentric residual region on chromosome 3 (Figure S2). Beyond these large contiguous blocks, we identified another 821 and 52 genes in the PT Mi21 families, respectively, that harbored high-confidence SNPs in the RNAseq data, yet were not contiguous with any of the large residual introgression regions. It is unlikely that there was sufficient recombination in the BC_5_ populations to generate these independent blocks; rather these genes likely have moved genomic coordinates in the landraces relative to B73, and actually reside inside one of the large introgression regions [30, 31]. However, none of these genes had genotypes that were perfectly correlated with genotypes at the *Inv4m* locus, so we excluded them all from further analysis.

**Fig 3.**
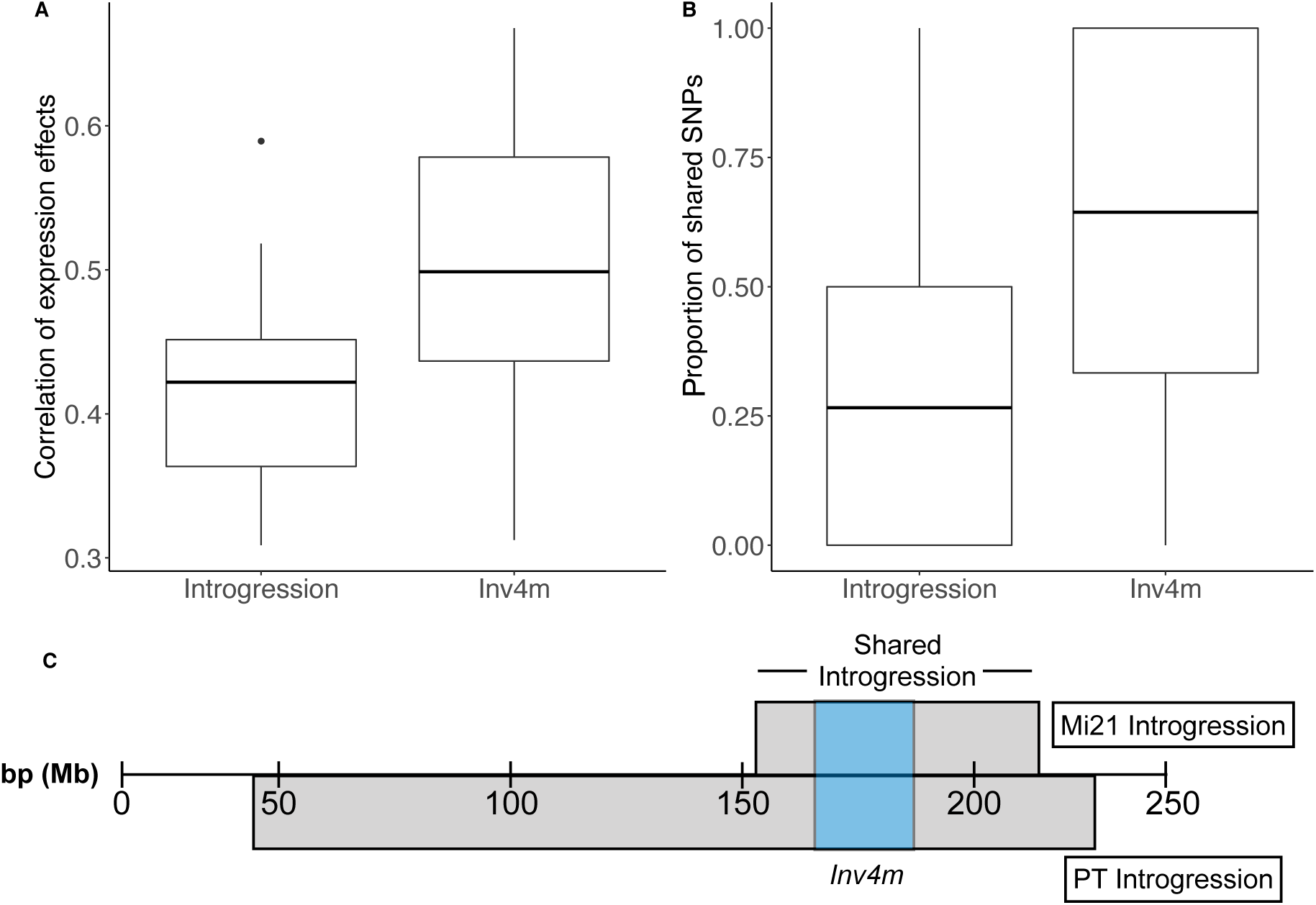
Sequence and expression divergence between landrace donors is greater outside of *Inv4m*. A) Boxplots show the correlation between expression effects of *Inv4m* genotype between the two donor populations for genes inside *Inv4m* or outside *Inv4m* but in the region of shared introgression. Correlations were measured separately in each of the 18 tissue:temperature combinations. B) Boxplots show the proportion of shared SNPs per gene for genes inside or outside *Inv4m*. C) Diagram of PT and Mi21 introgressions containing *Inv4m* on chromosome 4.

In the segregating families, there were a total of 7,236 genes from PT, and 4,095 genes from Mi21, 355 of which were within *Inv4m*. Of all genes from the landrace donor which were highly correlated with the *Inv4m* genotype (*ρ >* 0.5), 14% and 29% resided inside *Inv4m* in the PT and Mi21 families respectively. Therefore, many associations of traits with the *Inv4m* haplotype that we observe in one population may be caused by functional variants in regions flaking the inversion, rather than functional variants inside the inversion itself. However, based on our GWAS results, only variants actually inside the inversion (or in the breakpoint regions) are likely to be causal for the locus’s effect.

The proportion of shared SNPs between PT and Mi21 families within *Inv4m* was significantly higher than in the flanking introgressed regions (Fig 3B, t = −10.773, df = 357.8, p-value *<* 2.2e-16). The same pattern occurs for gene expression variation attributed to landrace genotypes for each gene: the landrace genotype’s effect on genes inside *Inv4m* are more correlated than landrace genotype effects in flanking regions (Fig 3C, t = −2.8869, df = 27.768, p-value=0.007) across the 18 tissue:temperature combinations in our experiment. Therefore, phenotypic effects associated with *Inv4m* that replicate across the two donor populations are likely caused by functional variation inside *Inv4m* rather than in residual landrace DNA present in each population (Fig 3). We also quantified the correlation between genetic and expression divergence between *Inv4m* sources, and found that they were positively correlated (r=0.79, p=2.2e-16, df=76, r^2^=0.6357).

### Effects of *Inv4m* on genome-wide gene expression

Of the 432 tissue samples collected (nine tissues *×* two temperature treatments *×* two segregating families *×* two *Inv4m* arrangements *×* three biological replicates *×* two experimental replicates), we excluded 53 samples with fewer than 100,000 reads, leaving a total of 379 samples. We detected a total of 23,428 unique genes present where at least a third of the samples had 10 or more counts in at least one of the tissue:treatment combinations (with an average of 17,016 genes per tissue) for a total of more than 306,000 gene expression traits. We visualized the overall transcriptome variation among samples using multi-dimensional scaling (Figure S3). Samples clustered predominantly by tissue, with an additional slight separation by temperature treatment, but no visible separation by family or genotype at the *Inv4m* locus. This was expected because only *∼* 7% of the genomes differed among samples. Among tissues, all leaf samples except the S2 leaf base formed one major cluster, the two root samples formed a second cluster, and the stem and SAM and S2 leaf base tissues formed separate individual clusters.

To assess the effects of highland alleles within *Inv4m* on global gene expression and plant development and physiology, we focused on genes with expression associated with *Inv4m* genotype that resided in “clean” genomic regions, so that the local genotype of each gene could not affect the association results. Overall, we identified 11,842 unique genes associated with genotype at *Inv4m* in the PT families and 12,482 genes in the Mi21 families, both using a 5% local false sign rate (*lfsr*) threshold for significance (Table 1). The number of associated genes varied by tissue and temperature treatment with a range of 1,932-5,753 and 1,018-6,000 genes identified per tissue in the PT and Mi21 families respectively. Of these genes, 285-1646 per tissue replicated across both donor families, where replication required *lfsr <* 5% and the effect in the same direction in both families in each tissue:temperature where the effect was significant. This reduced list of genes constitutes *Inv4m-regulated* candidate genes, and comprised 8 *−* 41% of the differentially expressed genes in the PT family, and 11 *−* 38% of the differentially expressed genes in the Mi21 family. Candidate *Inv4m-regulated* genes were distributed across the genome, with no visible clustering by chromosome.

In contrast, we detected only 38-413 genes per tissue in the PT family and 435-2398 genes per tissue in the Mi21 family with significant genotype-treatment interactions at a 5% *lfsr* threshold. Of these, 4-23 genes per tissue were shared across the two families (Table 2).

**Table 2.** Number of differentially expressed genes in each population and their overlap using a local false sign rate threshold of 5% across tissues and temperature treatments.

| Tissue | Cold |  |  | Warm |  |  | G × E |  |  |
| --- | --- | --- | --- | --- | --- | --- | --- | --- | --- |
|  | PT | Mi21 | Shared | PT | Mi21 | Shared | PT | Mi21 | Shared |
| V1 Leaf tissue | 2415 | 1534 | 285 | 2107 | 2330 | 453 | 105 | 575 | 6 |
| V1 Primary root | 1932 | 3516 | 495 | 3106 | 2759 | 649 | 84 | 1377 | 8 |
| V3 leaf base | 2550 | 3785 | 789 | 2809 | 3761 | 932 | 73 | 843 | 6 |
| V3 leaf sheath | 4047 | 5126 | 1646 | 3321 | 4049 | 1062 | 469 | 742 | 26 |
| V3 Primary root | 5753 | 3143 | 1206 | 2144 | 2039 | 413 | 42 | 455 | NA |
| V3 S2 leaf base | 3527 | 1018 | 268 | NA | 5665 | NA | NA | 2573 | NA |
| V3 S2 leaf tip | 2806 | 3054 | 721 | 4170 | 3190 | 1249 | 136 | 1527 | 19 |
| V3 Stem and SAM | 3118 | 6000 | 1269 | 2569 | 2961 | 752 | 190 | 1509 | 24 |
| V3 Leaf tip | 2207 | NA | NA | 4225 | 2453 | 997 | 362 | NA | NA |
| Totals | 12574 | 12485 | 2649 | 11468 | 12233(20.7%) | 2542 | 1306 | 5006 | 85 |

### *Inv4m-regulated* genes are enriched for diverse biological processes

We extracted the Gene Ontology (GO) terms associated with *Inv4m-regulated* genes and tested for enriched terms. For each tissue:treatment experiment, we separately tested for enrichments among the genes up-regulated, down-regulated, or both by *Inv4m*. We identified 596 enriched categories overall, with 0-152 categories enriched per tissue and temperature treatment at a 5% FDR. These GO terms provide candidate descriptors of the global effects of *Inv4m*. To reduce the number of categories found across the tissue:temperature treatments, we collapsed terms into clusters by semantic similarity and selected the most-enriched term across all tissue:temperature treatments in each cluster. After filtering for redundancy, we report twenty-two GO terms across the three main GO ontologies: two cellular component terms, two molecular function terms, and eighteen biological processes terms (Table 3).

**Table 3.**
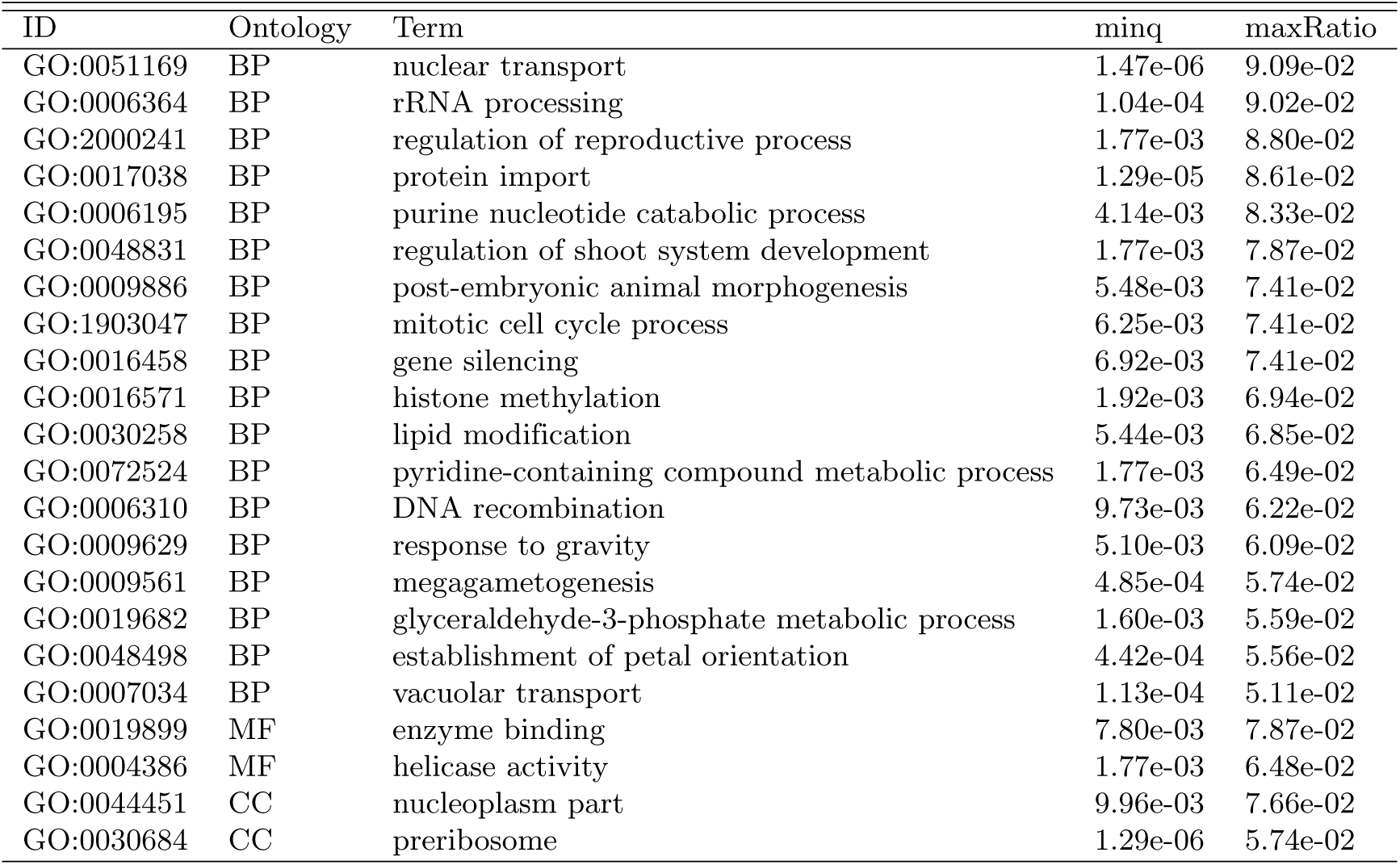
Gene ontology (GO) terms remaining after final filtering. A universal enrichment analysis was conducted on each tissue and temperature and directional (up-regulated, down-regulated, or both) combination for *Inv4m*-regulated genes. Terms were then ranked by enrichment score and grouped by a semantic similarity score of higher than 0.5. The top term in each semantic similarity group was then selected.

| ID | Ontology | Term | minq | maxRatio |
| --- | --- | --- | --- | --- |
| GO:0051169 | BP | nuclear transport | 1.47e-06 | 9.09e-02 |
| GO:0006364 | BP | rRNA processing | 1.04e-04 | 9.02e-02 |
| GO:2000241 | BP | regulation of reproductive process | 1.77e-03 | 8.80e-02 |
| GO:0017038 | BP | protein import | 1.29e-05 | 8.61e-02 |
| GO:0006195 | BP | purine nucleotide catabolic process | 4.14e-03 | 8.33e-02 |
| GO:0048831 | BP | regulation of shoot system development | 1.77e-03 | 7.87e-02 |
| GO:0009886 | BP | post-embryonic animal morphogenesis | 5.48e-03 | 7.41e-02 |
| GO:1903047 | BP | mitotic cell cycle process | 6.25e-03 | 7.41e-02 |
| GO:0016458 | BP | gene silencing | 6.92e-03 | 7.41e-02 |
| GO:0016571 | BP | histone methylation | 1.92e-03 | 6.94e-02 |
| GO:0030258 | BP | lipid modification | 5.44e-03 | 6.85e-02 |
| GO:0072524 | BP | pyridine-containing compound metabolic process | 1.77e-03 | 6.49e-02 |
| GO:0006310 | BP | DNA recombination | 9.73e-03 | 6.22e-02 |
| GO:0009629 | BP | response to gravity | 5.10e-03 | 6.09e-02 |
| GO:0009561 | BP | megagametogenesis | 4.85e-04 | 5.74e-02 |
| GO:0019682 | BP | glyceraldehyde-3-phosphate metabolic process | 1.60e-03 | 5.59e-02 |
| GO:0048498 | BP | establishment of petal orientation | 4.42e-04 | 5.56e-02 |
| GO:0007034 | BP | vacuolar transport | 1.13e-04 | 5.11e-02 |
| GO:0019899 | MF | enzyme binding | 7.80e-03 | 7.87e-02 |
| GO:0004386 | MF | helicase activity | 1.77e-03 | 6.48e-02 |
| GO:0044451 | CC | nucleoplasm part | 9.96e-03 | 7.66e-02 |
| GO:0030684 | CC | preribosome | 1.29e-06 | 5.74e-02 |

### Identifying candidate causal alleles within *Inv4m*

To identify candidate genes within *Inv4m*, we measured the *Inv4m* genotype effect (i.e. *cis* effect) on each of the annotated genes located inside the *Inv4m* locus, and the correlation between the expression of these *Inv4m*-genes and all *Inv4m-regulated* genes (a total of 4642 unique genes, with a range of 89-713 genes per tissue; Figure4). We repeated this analysis in each of the 18 tissue:temperature treatments. For *cis*-genotype effects to be counted, we required the effects to be significant (*lfsr <* 5%) and in the same direction between the two donor populations.

Overall, of 355 annotated genes within the boundaries of *Inv4m*, 224 were expressed in both donors. Of those, 155 were differentially expressed in the same direction in both donors in at least one treatment, and 89 of those were significantly correlated with at least one *Inv4m-regulated* gene located on other chromosomes used to measure the global *Inv4m effect*, even after accounting for the effect of *Inv4m* itself (Figure 4). Of these, 6 genes in particular stood out as being correlated with a large number (*>* 3%) of the reporter genes: Zm00001d051908, Zm00001d051998, Zm00001d052075, Zm00001d052079, Zm00001d052153, Zm00001d052259, and 9 were differentially expressed between *Inv4m* alleles in 90% or more of the tissue:temperature combinations. These were: Zm00001d051872, Zm00001d051882, Zm00001d051987, Zm00001d052051, Zm00001d052136, Zm00001d052210, Zm00001d052242, Zm00001d052245, Zm00001d052269. Finally, three of the 89 candidate *Inv4m* genes were transcription factors: Zm00001d051879, Zm00001d052180, Zm00001d052229. These genes are reported in Table S2, with associated description, GO term(s), and previous GWAS trait associations.

**Fig 4.**
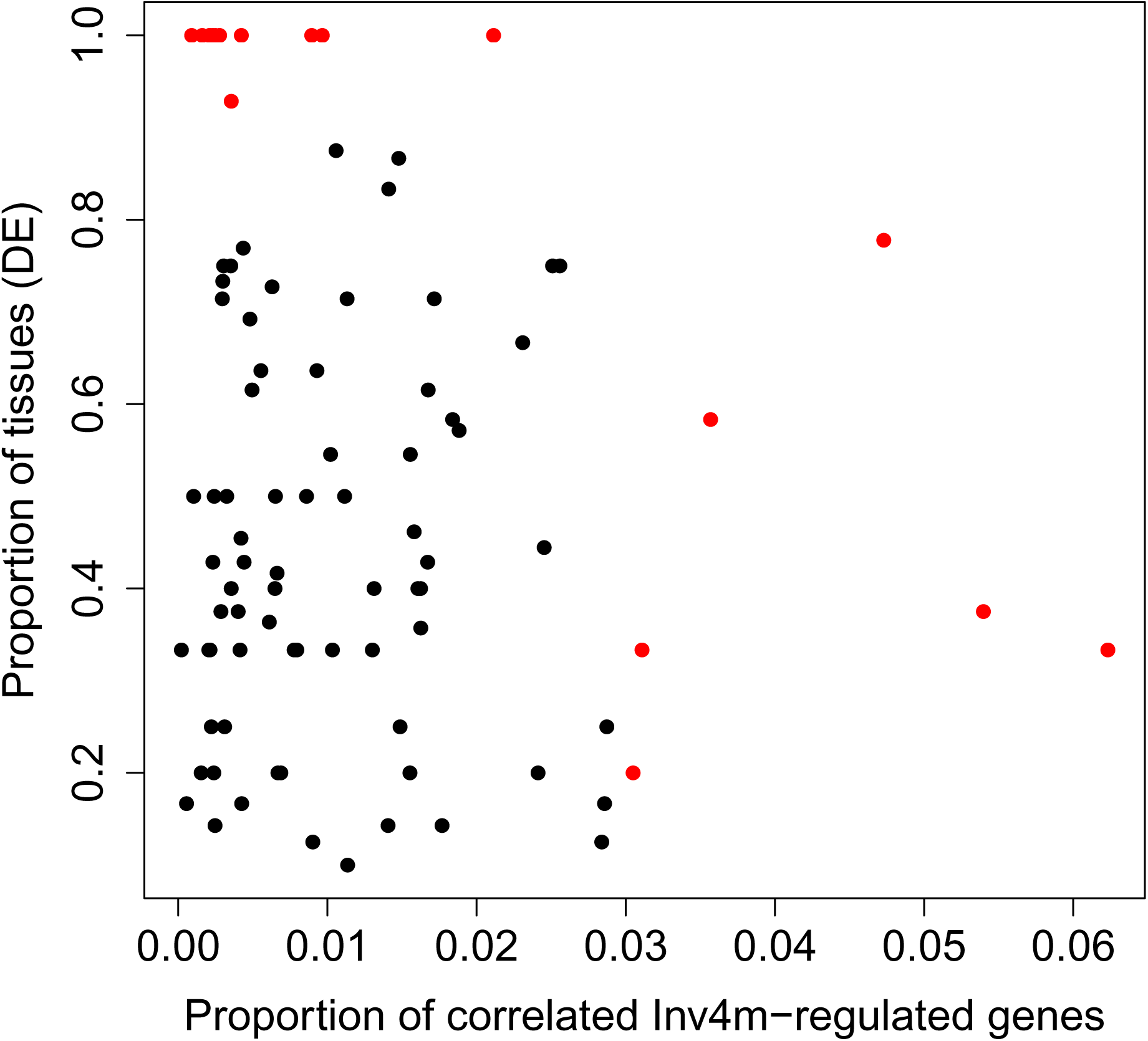
Scatter plot of candidate gene scans within *Inv4m*. Each point represents one of the 89 genes inside *Inv4m* that were both differentially expression by *Inv4m* genotype in both populations and correlated with at least one *Inv4m-regulated* gene. The x-axis represents the proportion of *Inv4m-regulated* genes correlated with the *Inv4m*-gene (p-value*<* 0.05). The y-axis represents the proportion of tissues in which the *Inv4m*-gene is differentially expressed (lfsr*<* 0.05 in both populations, and regulated in the same direction in both populations.)

Since the phenotypic association of *Inv4m* with flowering time is so large, we tested specifically for effects of the locus on the expression of genes with known roles in regulating flowering. Of the list of 48 flowering genes in Dong et. al. 2012 [32], 5 were consistently regulated by *Inv4m* in both populations in the same direction (Table S3). These expression effects were detected in the V1 and V3 primary root, S2 leaf tip, and stem and SAM tissues.

Another *a priori* candidate gene for the effect of the inversion is the microRNA MIR172c, which is located inside *Inv4m* at coordinates: 4:174154928-174155050. MIR172 has known roles in regulating developmental transitions as well the development of pigmentation and macrohairs in maize [33]. The expression of the pre-miRNA 172c was too low to assess differential expression in all but the leaf sheath tissue, and was not associated with *Inv4m* genotype in that tissue. However, of 31 genes that are predicted targets of miR172, 9 were consistently regulated by *Inv4m* in both populations in one-or-more tissues. Of these, six were down-regulated by the High haplotype of *Inv4m*. These genes and their associated descriptions are found in Table S4.

### De novo transcriptome assembly to search for novel genes

Finally, we used the RNAseqreads that did not map to the B73 genome to search for genes that may be present in the High haplotype of *Inv4m* but not in the Low haplotype, and thus may have been missed by our analysis. We assembled all un-mapped reads from samples carrying the PT or Mi21 haplotype at *Inv4m* using Trinity, and searched for de-novo transcripts with evidence for expression only in the samples carrying the High haplotype. We found 772 candidate transcripts. However, all were very lowly expressed. Only five had estimated transcript counts of at least 10 summed across all *PT* or *Mi21* samples, and none had estimated transcript counts of at least 10 in two or more samples per family. Therefore, we found no evidence that the High haplotype of *Inv4m* carries high-expressed genes that are not present in the Low haplotype.

## Discussion

We combined three methodological approaches to study the function of the chromosomal inversion, *Inv4m*, found in highland populations of domesticated maize (*Zea mays* ssp. *mays*). First, we used a large maize diversity panel designed for Genome-wide association studies (GWAS) to identify agronomic traits associated with *Inv4m*. Second, we created a two-donor panel consisting of two sets of segregating families segregating for both arrangements of the *∼* 13*Mb* inversion and grew them reciprocally in two temperature treatments to measure gene expression effects of *Inv4m* across nine tissues from early developmental plants. In total, we inspected *Inv4m* effects on more than 300,000 traits and employed statistical methods that leveraged the replication of the effects across conditions [34] to broadly characterize the effects of this locus. Finally, we inspected cis-regulatory variation on genes within the inversion to identify candidate genes responsible for the *Inv4m* effects.

### 0.1 Association analysis of inversion loci

GWASs in diversity panels and QTL mapping in biparental populations or NILs are complimentary methods for locating genetic loci associated with phenotypic variation. GWASs can pinpoint the location of functional variants more precisely than bi-parental mapping populations because they use diverse association panels that encompass a large number of ancestral recombination events [35]. However, GWAS approaches suffer from confounds due to population structure and non-equal relatedness leading to high frequencies of false positives [36]. QTL mapping populations, on the other hand, are limited in resolution due to the requirement for new recombination events between candidate loci.

Multi-parent populations like NAM [37] or MAGIC [38, 39] capture advantages of both methods, approaching the resolution of GWAS without the confounds of population structure. Our segregating families shared a common parent (B73), and thus are similar to a two-population NAM panel. NAM-QTL mapping can have high precision when the donor parents share the same alleles at a causal locus and do not share alleles at other loci. In our case, the two donor parents PT and Mi21 likely share many alleles other than *Inv4m* due to shared highland ancestry throughout the genome, so families from both parents will overlap at other causal alleles linked to *Inv4m*. However, our results show that the genetic similarity between PT and Mi21 is much higher inside *Inv4m* than in the remaining *∼*50Mb of shared residual landrace genome between the two donors. Therefore, while we cannot unambiguously attribute shared phenotypic effects across the two donors to *Inv4m*, the majority of effects of *Inv4m* that we identified are likely caused by functional variation within this locus itself, rather than due to other shared alleles between the two donors.

Our GWAS study of diverse maize landraces confirmed earlier results [26, 29] that highland haplotypes of *Inv4m* are strongly associated with agronomically important phenotypes including flowering time, the Anthesis-Silking interval, and several measures of yield. In each case, the effect of the highland haplotype was beneficial when grown in highland environments, but detrimental when grown in lowland environments, consistent with this single locus causing antagonistic pleiotropy [40] across elevation environments. This may explain the strong evidence for selection at this locus: its divergence in allele frequencies across elevations and the evidence that this locus introgressed into highland maize from the wild relative *mexicana* [23]. That the locus appears to independently control both flowering and yield traits is consistent with the idea that *Inv4m* contains multiple important variants that each contribute to phenotypic differences between lowland and highland maize [26, 29]. Because they are inherited as a single unit, inversions are thought to contribute to the prevalence of local adaptation despite high gene flow [41–43] by linking these multiple variants into a single *supergene* [44].

However, while compelling, due to the strong population structure in the diversity panel used above, we caution that the associations of *Inv4m* with the agronomic traits is still preliminary. *Inv4m* genotypes are highly correlated to overall genetic ancestry (as measured by PC1 calculated using all other chromosomes except chromosome 4) within Mexico. We corrected for ancesetry using genome-wide kinship (again excluding chromosome 4), but if this correction was incomplete, it may have lead to false-positive associations of phenotypic traits with *Inv4m*. An alternative explanation for the phenotypic associations above is that each trait has a polygenic basis, with small-effect loci distributed throughout the genome, each of which has subtle allele-frequency differences across elevations. We conducted a preliminary analysis of the PT and Mi21 segregating families in high elevation fields, but we did not have sufficient replication to detect any effect. We are following up with experiments in highland field conditions to test the effects of the inversion on gene expression and fitness.

### 0.2 Progress towards inversion fine-mapping

Whether or not *Inv4m* directly controls flowering time and yield, neither GWAS not NAM-QTL mapping themselves can provide direct insight into which functional variant(s) *inside* the locus are responsible for these phenotypic effects. And unlike a typical GWAS peak that may cover dozens of possible variants in high LD with a tested marker, there may be hundreds of thousands of variants between the two haplotypes of *Inv4m* within this *∼* 13Mb region, any of which could be responsible for some of the phenotypic effect of the locus. Neither GWAS nor QTL mapping itself can prioritize any of these variants, because they remain in near-perfect LD in either type of panel.

Ultimately, identifying specific functional loci within *Inv4m* will require experimental mutagenesis or other genetic perturbations within the locus. However, we aimed to begin to characterize the diversity of functional variants in this locus using gene expression analysis. We used gene expression in four distinct ways to dissect the functional variation captured by *Inv4m*: 1) by analyzing genome-wide gene expression responses to *Inv4m* to mine for phenotypic effects across *>*300,000 traits; 2) by analyzing local *cis*-regulatory effects of the locus on the 355 genes within *Inv4m*; 3) by analyzing the co-expression between *Inv4m* genes and the rest of the genome; and 4) by assemblying un-mapped transcripts into de-novo gene models. The first analysis provided an exceptionally detailed phenotypic dissection of the total effects of the *Inv4m*, and showed that there are likely many distinct components to the cellular and physiological effects of *Inv4m*. The second analysis provided an estimate of the density of functional variants within the *Inv4m* locus: we detected likely *cis*-regulatory variation affecting 155 genes. The third analysis showed that many of these cis-regulatory variants may have functional consequences beyond the immediate genes they regulate. The fourth analysis did not give any compelling results, but may be useful in other systems.

By studying gene expression effects of *Inv4m* in two relevant environmental contexts (hot and cool temperatures) and across nine distinct tissues, we aimed to maximize our ability to discover developmental and phenotypic effects of the locus. It is certainly possible that we missed important phenotypic effects of *Inv4m* by sampling only tissues on young plants - effects on pathways specific to reproductive tissues were likely missed. However, we selected the nine tissues based on the published maize gene expression atlas [45] so as to capture as much variation as possible in expression profiles, given the experimental constraints on how large we could let the plants grow in our growth chambers.

Overall, our expression results show that *Inv4m* affects many disparate biological processes in young maize tissues. The strongest Gene Ontology enrichment signals among *Inv4m*-regulated genes were in terms related to mRNA and protein processing around the nucleus (nuclear transport and import, and the pre-ribosome). We also found evidence of effects on epigenetic regulation, cell-cycle processes, metabolism, and development. None of these results provide clear explanations for the effects of *Inv4m* on flowering time and yield. However both flowering and yield are highly complex traits that are affected by many aspects of development, physiology, and stress responses, and so the mechanistic links among these traits may not be obvious [37]. We looked more specifically at *a priori* candidate genes for flowering and yield traits both inside and outside *Inv4m* and found possible effects on several of these genes, but no strong enrichment of *Inv4m* effects on either class.

Among the genes within *Inv4m*, nearly 70% of those expressed high enough to measure showed evidence of *cis*-regulatory variation among alleles. While some of these genes may share regulatory elements, its likely that the majority of these genes are affected by independent genetic variants. This suggests that the two haplotypes of *Inv4m* harbor a large number of functionally relevant genetic differences. However, does *Inv4m* harbor more functional variants than any other similarly sized introgression among maize landraces? To test this, we compared the number of genes (genome-wide) correlated with *Inv4m* in each segregating family to the number of genes that show similar expression in both donor families. The latter genes are those we believe are truly affected by *Inv4m*, while the remainder are likely regulated by PT or Mi21 alleles that reside in introgressed genomic outside of *Inv4m* in each population. In both populations, the proportion of *Inv4m* candidates among all *Inv4m*-correlated genes is roughly similar to the relative sizes of *Inv4m* to the whole chromosome 4 introgression in each population. This suggests that introgressing any region from PT or Mi21 into B73 will cause diverse effects on gene expression, and that *Inv4m* is not exceptional in the magnitude of these perturbations.

Together, these results imply that the *Inv4m* locus has many effects on corn development and physiology, and therefore it’s contribution to local adaptation is complex and not simply a change to major flowering or yield-related genes. The incorporation of *∼* 13Mb of the genome of *mexicana* likely brought with it a large number of functional variants that have both positive and negative effects on many molecular traits, most of which are not macroscopically visible, but may still impact performance in different conditions.

## Conclusions

This study represents a broad characterization of an adaptive chromosomal inversion. Our results give insight into the role of this inversion in adapting to high altitude environments. GWAS results show that *Inv4m* is associated with faster flowering and higher yield in highland common gardens. The molecular roles of genes within the inversion are summarised by the phenotypic effects of *Inv4m*-regulated genes (Table 2) and enriched GO terms (Table 3), and the candidate gene set within *Inv4m* (Table S2). Fine-mapping in this region is required to further dissect the functional role of loci within *Inv4m*, but will have additional challenges due to suppressed recombination between heterokaryotypes. Novel genomic technologies, such as a CRISPR/CAS system [46] that can reverse the orientation of the High *Inv4m* haplotype could be used to induce recombination across the newly collinear genomic regions, allowing the localization of specific effects of the different variants linked in this locus.

## Materials and methods

### Population Genetics of *Inv4m*

We downloaded unimputed genotype-by-sequencing (GBS) data from 94,726 loci on chromosome 4 for 4,845 maize plants from the SeeD-maize GWAS panel [26, 47] and ran a principal components analysis on all positions within the *Inv4m* locus (between 168832447 and 182596678 in AGPv2 coordinates, [24]) with *<* 25% missing data and minor allele frequencies *>* 0.05. PC1 explained 22% of the genetic variation among these plants in this interval. Scores on PC1 neatly divided plants into three groups, representing the two homozygous classes at *Inv4m* and their heterozygotes. We cross-referenced plants with landrace passport data from Germinate 3, the CIMMYT Maize Germinate Database germinate.cimmyt.org/maize/ and extracted country of origin, latitude, longitude and elevation records. All but 7 of the plants containing the minor haplotype at *Inv4m* were from Mexico, so to study associations with elevation and to calculate other diversity statistics, we subsetted to only those plants collected in Mexico.

To calculate the association of *Inv4m* with elevation, we divided landraces into 100m bins, calculated the haplotype frequency of the minor haplotype in each bin, and fit a loess curve to the log-transformed haplotype frequencies, weighted by the number of landraces in each bin. Based on this analysis, we labeled the minor haplotype in Mexico “High”, and the major haplotype “Low”. We used the R function *HWExact* from the *HardyWeinberg R* package to test genotype counts against the Hardy-Weinberg expectation. We calculated diversity statistics *π* and *θ* separately for plants homozygous for the “High” or “Low” haplotypes at *Inv4m* across all of chromosome 4 in 500 marker windows using TASSEL 5 [48]. Because many more plants were homozygous for the “Low” haplotype, we randomly sampled 371 landraces to calculate the diversity statistics to make the sample sizes equal.

### Association of *Inv4m* with agronomic traits

We re-analyzed phenotypic data from the F1 Association Mapping (FOAM) panel of Romero-Navarro *et al* [26] and Gates *et al* [29] to more fully characterize associations signatures of *Inv4m*. Full descriptions of this experiment and data access are described in those references. We downloaded BLUPs for each trait and line from Germinate 3, and subsetted to only those lines with GBS genotype data from Mexico. We fit a similar model to the GWAS model used by Gates *et al* [29] to estimate the effect of *Inv4m* genotype on the trait’s intercept and slope on trial elevation, accounting for effects of tester ID in each field and genetic background and family effects on the trait intercept and slope using four independent random effects. We implemented this model in the *R* package *GridLMM* [49]. We extracted effect sizes and covariances conditional on the REML variance component estimates and used these to calculate standard errors for the total *Inv4m* effect as a function of elevation. To test whether the phenotypic effects of *Inv4m* on yield components could be explained as indirect effects via flowering time, we additionally re-fit each model using Days-To-Anthesis as a covariate with an independent effect in each trial.

### Experimental material for isolating *Inv4m*

To directly assess phenotypic effects of the *Inv4m* locus, we selected two highland landrace accessions which both carry the High haplotype of *Inv4m*, Palomero Toluqueno (PT) and an accession from the Cónico landrace. These *Inv4m* landrace accessions were obtained through the International Maize and Wheat Improvement Center (CIMMYT); PT came from accession mexi5 (referred to henceforth as PT) and Cónico from accession Michoacán 21 (referred to henceforth as Mi21). B73 is a modern inbred from the United States that carries the non-inverted haplotype at the *Inv4m* locus. Both landraces were crossed with B73 and one resulting F1 individual from each cross was backcrossed to B73 for 5 generations, selecting on a diagnostic SNP for *Inv4m* each cycle with a cleaved amplified polymorphic sequence (CAPS) assay. The diagnostic SNP is at position 4:179617762 (Matt Hufford, Personal communication). DNA was extracted from leaf tissue using a Urea lysis buffer extraction protocol (https://github.com/RILAB/lab-docs/wiki/Wetlab-Protocols). Primers were designed to amplify the fragment of DNA carrying the diagnostic SNP (Forward: CTGAGCAGGAGATGATGGCCACTC; Reverse: GGAAAGGACATAAAAGAAAGGTGCA). Amplification consisted of 5 minute denaturation at 95*^◦^*C, 35 cycles of 95-60-72*^◦^*C for 30 seconds each, 7 minutes of final extension step at 72*^◦^*C, followed by a 4*^◦^*C hold. Amplified DNA was then digested with the *Hinf 1* enzyme for 1 hour at 37*^◦^*C, and the resulting product was run out on a 1% agarose gel for genotyping.

Two of the resulting BC_5_ individuals identified to be heterozygous for *Inv4m* were self-pollinated per population to produce BC_5_S_1_ families segregating for *Inv4m*.

### Reciprocal transplant experiment to identify phenotypic effects of *Inv4m*

We planted seeds from the four segregating families (two parents per *Inv4m* donor) in the UC Davis controlled environment facility growth chambers. Chambers were programmed to mimic temperatures in Mexican lowlands (22*^◦^*C night, 32*^◦^*C day, 12 hr light) and highlands (11*^◦^*C night, 22*^◦^*C day, 12 hr light). Kernels were soaked in distilled H_2_0 for 12 hours and planted in 10.2cm x 34.3cm nursery pots (Steuwe & Sons: CP413CH) in a substrate mixture composed of a 3:1 ratio of Sungro Sunshine Mix #1 to sand. Pots were organized in racks with 9 pots per rack (Steuwe & Sons: tray10). Plants were watered every other day with a 1x Hoagland nutrient solution, and emergence was recorded daily. The experiment was replicated and growth chambers were switched to account for variation between instruments between replicates (See Figure S1 for a graphical workflow). The first replicate of the experiment began March, 2017 and the second replicate began April, 2017.

Two seeds were planted in each pot, one in the center and one near the corner, and a total of nine tissues were sampled from the two plants when they reached specific developmental stages. These nine tissues were selected to maximize the diversity of gene expression profiles based on the transcription atlas of [45]. Corner plants were removed from the pot and sampled when they reached the V1 stage, while center plants were sampled when they reached the V3 stage. Two tissue types were sampled from the V1 stage, and 7 tissue types were sampled from the V3 stage (Table 1). Sampling occurred between 2 and 4 hours after simulated sunrise. Plant tissue was placed in 2 ml centrifuge tubes, immediately flash frozen in liquid nitrogen, and stored at −70*^◦^*C.

### *Inv4m* Genotyping and RNA sequencing

We used the same DNA extraction and CAPs genotyping methods as previously described to genotype the seedlings for the *Inv4m* haplotype. We harvested and successfully genotyped 364 individual plants from families segregating for *Inv4m*. The Low, Heterozygote, and High *Inv4m* haplotypes were segregating in the PT families at a 42:78:45 ratio, within Hardy-Weinberg equilibrium (HWE; D= −2.24, p-value=0.53). The inversion was segregating in the Mi21 families at a 59:97:43 ratio, also within HWE (D= −0.928392, p-value=0.7768964).

We randomly sampled 3 biological replicates per experimental replicated from each tissue and temperature treatment for the two homozygous *Inv4m* genotypes (Mi21 & PT), a total of 432 samples from 96 plants. Approximately 20 mg of tissue from each sample was placed in a 2ml centrifuge tube, flash-frozen in liquid nitrogen and ground using stainless steel beads in a SPEX Geno/Grinder (Metuchen, NJ, USA). mRNA was extracted using oligo (dT)_25_ beads (DYNABEADS direct) to isolate polyadenylated mRNA using the double-elution protocol. We prepared randomly primed, strand specific, mRNA-seq libraries using the BRaD-seq [50] protocol with 14 PCR cycles. Samples underwent a single carboxyl bead clean-up, were quantified using the Quant-iT^TM^ PicoGreen dsDNA kit, and then normalized. We took 2ng per library and multiplexed 96 samples for sequencing. Each multiplexed library was sequenced on 1 lane of a Illumina HiSeq X platform, generating a mean of 4,241,500 reads per sample. Raw reads were quality checked using FastQC v.0.11.5 [51]. Adapter sequences, low quality reads (q*<*20), and sequences less than 25 bp were removed using Trimmomatic v.0.36 [52].

### Effects of genotype at *Inv4m* on seedling emergence

The effect of *Inv4m* on seedling emergence were analyzed using the following random slope and intercept model for each donor and temperature treatment separately:

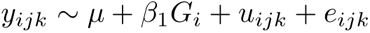

*y_ijk_* is the emergence time for individual plant *k* in experimental replicate *j* in *Inv4m* genotype *i*. *µ* is the model intercept, *β*_1_*G_i_* is the effect of *Inv4m* genotype, **u** = **u_ijk_** is a random effects term for the experimental replicate, and *e_ijkl_* is residual error. Variance components, coefficients and standard errors were estimated by REML using the *R* function *lmer* [53], and p-values were calculated using conditional F-tests [54].

### Population characterization

BC_5_S_1_ plants are expected to contain *∼* 3% residual DNA inherited from the donor parent across the remaining 9 chromosomes. We used the RNAseq reads from each plant to genotype all residual regions across the genome by calling variants in the expressed regions. Paired reads that passed filtering were aligned to the B73 reference genome version 4 [55] using hisat2 [56], and variant loci were called using GATKv3 [57, 58]. We ran `MarkDuplicates, SplitNCigarReads and HaplotypeCaller` on every sample, including all 435 samples from our segregating families and an additional 46 B73, 48 PT and 47 B73-PT F1 samples from plants run in parallel with the families but were not otherwise used in this experiment, and then ran `GenotypeGVCFs` on all samples jointly. We next used `SelectVariants` to extract SNPs, and `VariantFiltration` to remove SNPs with FS score *<* 30 and QD *>* 2.0. We further filtered for SNPs called homozygous-reference in all B73 samples, and which exhibited allele frequencies *>* 1/8 and *<* 7/8 in either the PT or Mi21 families (expected frequencies of each variant should be 0.25 or 0.5 depending on recombination between the two BC_5_ parents of each population, but we allowed for some sampling error). We used this set of highly filtered SNPs for each population to genotype each of the individual plants. For each plant, at each locus we first combined all genotype likelihoods across all RNA samples from the same sample. We then identified the approximate breakpoints of the introgressed regions by inspecting the density of variant sites. We identified 3 regions (on chromosomes 2, 4 and 5) in the PT family and 2 regions in the Mi21 family (on chromosomes 3 and 4). Within these introgressed regions, we used `R/QTL` [59] to assign genotype probabilities across the *Inv4m* locus for each plant, allowing `error.prob =` 0.2. Finally, we observed that several genes outside these 5 introgressed regions each of which exhibited *>*= 2 SNPs relative to the reference. We hypothesized that these genes may have a different chromosome location in the landraces relative to B73, and actually reside inside one of the 5 introgressed regions. We therefore assigned their genotype to the most common genotype among these variant loci.

### RNA quantification

To quantify gene expression, we ran `kallisto v.0.42.3` [60] separately on each sample using the B73 AGPv4.36 transcript models downloaded from the maize genome database [55]. We limited to only one bootstrap replicate, and then summed the transcript counts for each gene. Genes were retained for analysis when at least a third of the samples had 10 or more reads. Gene counts were normalized using the weighted trimmed mean of M-values (TMM) with the *calcNormFactors* function in *edgeR* [61]. Normalization using TMM reduces bias of very highly and lowly expressed genes. The *voom* function [62] in the *limma* package [63] was used to convert normalized reads to log2-counts per million (log2CPM), estimate a mean-variance relationship, and assign each observation a weight based on its predicted variance. Observation-weights were used in downstream analyses to account for heteroscedasticity. We estimated batch effects using the *removeBatchEffects* function in limma using the experimental replicate as batch, which corrected the log2CPM expression values. Global patterns of gene expression across the experiment were visualized with the *plotMDS* function from *edgeR*.

### Analysis of *Inv4m* effects on gene expression

We divided genes into three groups to estimate the effects of *Inv4m* or other introgressed landrace alleles, based on whether each gene resided in a “clean” genomic region with only B73’s allele present in the families, inside the *Inv4m* locus itself, or if it resided within one of the genomic blocks containing any of the residual landrace genome outside of *Inv4m*. Each group of genes served a different purpose in the analysis of *Inv4m*. Genes in the “clean” region were used to assess the effects of *Inv4m* on global gene expression and indirectly assess the effects on development and physiology more broadly. Genes inside the *Inv4m* locus were scanned for candidate alleles underlying *Inv4m’s* effects. Genes in the residual introgression blocks were used as controls to assess the similarity of PT and Mi21 alleles in other genomic loci, as well as compare effect size and expression correlation with *Inv4m*.

For genes that resided in “clean” genomic regions with only B73’s allele present in the segregating families (approximately 89.8% of genes expressed in both donors), we estimated the effect of the *Inv4m* locus separately in each *Inv4m* donor, temperature treatment, and tissue using the linear model:

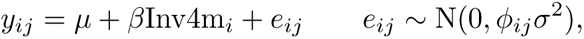

where *y_ij_* is a normalized, batch-corrected log2CPM value for a single gene in a single sample, *µ* is the intercept for that gene in the particular population, environment and tissue, *β* is the corresponding effect of the landrace *Inv4m* haplotype, and *e_ij_* is the model residual, which is assumed to be independent of all other residuals and have variance proportion to *φ_ij_*, the empirical weight factor calculated by `voom`. We fit this model to the whole set of “clean” genes using the *lmFit* function from *limma*, and extracted *β* and its standard error (*|β|/t*). We leveraged the correlations in effect sizes across tissues and environments to improve effect size estimates and identify a union set of genes regulated by *Inv4m* by combining results across tissues and environments using the *mash* method [34] implemented in the *mashr* R package. *mash* was run separately for the two segregating families.

We also fit a separate model to test for interactions between *Inv4m* and the temperature environment, separately for each *Inv4m* donor and tissue:

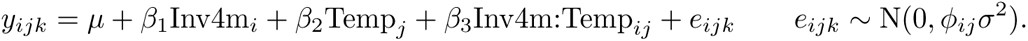

This model adds *β*_2_, the main effect of temperature environment on expression, and *β*_3_, the interaction between *Inv4m* and temperature. However it is less flexible than the first model because the residual variance *σ*^2^ is constrained to be equal for the two temperature environments. This model was also fit to each gene using *lmFit*, and the estimate of *β*_3_ and its standard error were extracted. We again used *mashr* to identify the union set of genes affected by this interaction.

For genes residing inside the *Inv4m* locus, we fit the same two statistical models with the *lmFit* function (both sets of genes were analyzed jointly to leverage the empirical bayes shrinkage of standard errors). However, we did not include these genes in the multiple adaptive shrinkage analysis.

For genes residing outside *Inv4m*, but within one of the genomic blocks containing residual landrace DNA in both donors, we fit a slightly different statistical model:

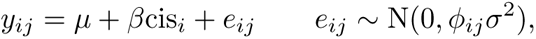

where *cis* is the local genotype of the gene, and *β* is the associated effect. For genes in residual genomic blocks on chromosome 4, the *cis* genotypes were highly correlated with the *Inv4m*, so some of the *cis* effect may have been caused by *Inv4m*, but these effects were difficult to separate statistically. However for genes on other chromosomes, the two genotypes were largely uncorrelated.

### Sequence divergence, and expression correlation between genes within the High *Inv4m* haplotype and the residual genomic regions

We estimated the sequence divergence between the two landrace donors and B73 for each gene within the chromosome 4 introgression containing *Inv4m* present in both families. For each gene, we calculated the genetic similarity of the PT and Mi21 alleles relative to B73 by counting the number of shared SNPs divided by total number of observed SNPs within each gene window, using only the highly filtered SNP set described above. We calculated the correlation in effect sizes between donors for each tissue and temperature combination. T-tests were used to determine whether genetic similarity and expression correlation were higher within *Inv4m* relative to the shared introgressed region.

### Gene ontology enrichment

Genes were assigned to gene ontology (GO) categories for functional annotation using an updated ontology annotation [64] which we expanded to include all ancestral terms for each gene with the *buildGOmap* function of the R package clusterProfiler [65]. Genes in “clean” genomic regions which responded to the High *Inv4m* haplotype in the same direction in both donor populations were classified as *Inv4m-regulated* and tested as foreground genes in a GO enrichment analysis. Genes that were expressed in each tissue and temperature combination in both donor populations, but were not *Inv4m-regulated* were included in the set of background genes. We calculated the enrichment of each GO term using the *enricher* function in the clusterProfiler R package. We selected GO terms with a false discovery rate less than 1% after a Benjamini-Hochberg multiple test correction. We then calculated the percent of genes in each GO terms that were *Inv4m-regulated*, and ranked all GO terms by their maximum enrichment across conditions. We then selected the highest enriched GO term among each set of GO terms that had a semantic-similarity *>*0.5 as the representative term set.

### Candidate gene pathway assessment among *Inv4m-regulated* genes

We inspected two additional candidate gene sets: genes known to regulate flowering time in maize [32], and genes regulated by the microRNA miR172. *Inv4m* has previously been associated with flowering [26]. miR172 is a highly conserved micro-RNA across the plant kingdom that regulates development and flowering time and one miR172 gene, *miR172c* is located inside *Inv4m*. For miR172 targets, we found the mature sequence for zma-miR172c: “AGAAUCUUGAUGAUGCUGCA” from miRBase http://www.mirbase.org, and used this as a query of the Plant Small RNA Target Analysis Server (psRNATarget, http://plantgrn.noble.org/psRNATarget), and collected all predicted target genes. We also used TAPIR’s pre-computed target genes for *zma-miR172a-b-c-d*. These two categories of genes were inspected by hand for evidence of regulation by *Inv4m*.

### Candidate genes within *Inv4m*

We used the intersection of two separate methods to identify candidate adaptive genes within *Inv4m*. First, we quantified the proportion of conditions (tissue:temperature combinations) that each gene within *Inv4m* was differentially expressed according to *Inv4m* genotype. To be considered differentially expressed, the gene needed to be differentially expressed in both *Inv4m* donor populations and in the same direction. Genes where at least one donor was not expressed were removed from this analysis per condition. As a complimentary approach, we calculated the correlation in expression between each *Inv4m* gene and all the genes in “clean” genomic regions that we determined to be *Inv4m-regulated* above. We then quantified the proportion of these correlations that were significant and in the same direction in both donor populations for each gene. For this analysis, we used the *lm* function in R to implement a linear model with the *Inv4m-regulated* gene expression as the response variable, and the *Inv4m* gene’s expression and *Inv4m* genotype as predictors.

### De-novo assembly of novel genes

We collected all un-mapped reads from the samples homozygous for the high *Inv4m* haplotype, and used Trinity v2.4 [66] to assemble un-annotated transcripts using default settings. We then used Kallisto to quantify the expression of each of these novel transcripts using the un-mapped reads from each RNAseq sample. To search for candidate “novel” genes in the highland haplotype, we filtered for Trinity genes that had zero estimated counts in any of the samples that were homozygous for the B73 haplotype of *Inv4m*, but had non-zero estimated counts in at least 2 samples homozygous for the PT or Mi21 haplotype in each segregating donor family (to exclude genes that may reside in either PT or Mi21).

## Supporting information

**S1 Figure.** Graphical workflow of breeding design, growth chamber experiment, sampling, sequencing, and analytical pipeline.

**S2 Figure.** Genotyping results using RNAseqdata, and genotype correlation with *Inv4m*.

**S3 Figure.** Multidimensional scaling plot of normalized and batch corrected gene expression (log2CPM) of the 500 highest expressed genes across our dataset using the *plotMDS* function. The plot represents a two-dimensional log2 fold change distance between each sample colored in three different ways to display how each factor effects the structure of the data. A) tissue B) *Inv4m* genotype and C) Temperature.

**S1 Table.** Summary of linear mixed effects model of seedling emergence for each *Inv4m* donor and temperature treatment

**S2 Table.** Candidate genes within the High *Inv4m* haplotype. The List column identifies how genes got on the list from the original list of 89. Correlated trans means that they are correlated with more than 3% of genes that were *Inv4m-regulated*, consistent DE are genes that were differentially expressed in more than 90% of the tissue:trans conditions that they were expressed in, and TF=transcription factor.

**S3 Table.** Flowering Genes from a priori list (Dong et. al. 2012) that were *Inv4m-regulated*.

**S4 Table.** miRNA172 target genes that were *Inv4m-regulated*. Homologous gene descriptions are from Arabidopsis thaliana database (https://www.arabidopsis.org/). GO annotation descriptions are the biological function (http://maizemine.rnet.missouri.edu/). TF=Transcription factor.

## Acknowledgments

We would like to thank Brittney Gillespie, Luis Avila, and Po-Kai Huang for help sampling tissue, and to the staff at the controlled environment facility for helping us set up and maintain light and temperature regimes in the growth chambers. This work was supported by National Science Foundation (Grant No. 1546719).

