## Supplemental for "Gene regulatory effects of a large chromosomal inversion in highland maize" (from another version of this preprint: version 1, 2019-12-02)

### Supplemental Figures and Tables

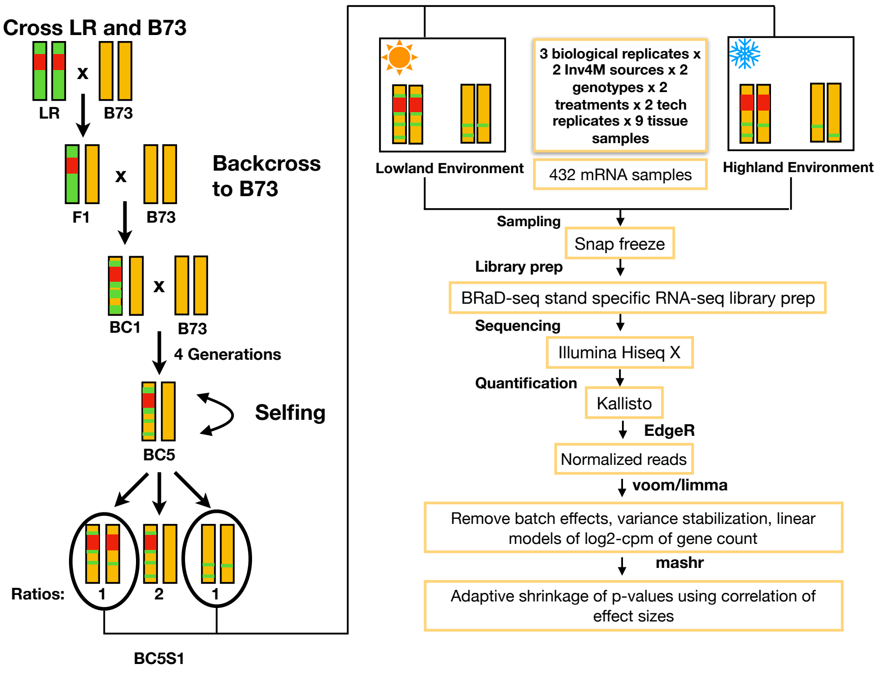


Figure 1: Graphical workflow of breeding design, growth chamber experiment, sampling, sequencing, and analytical pipeline.

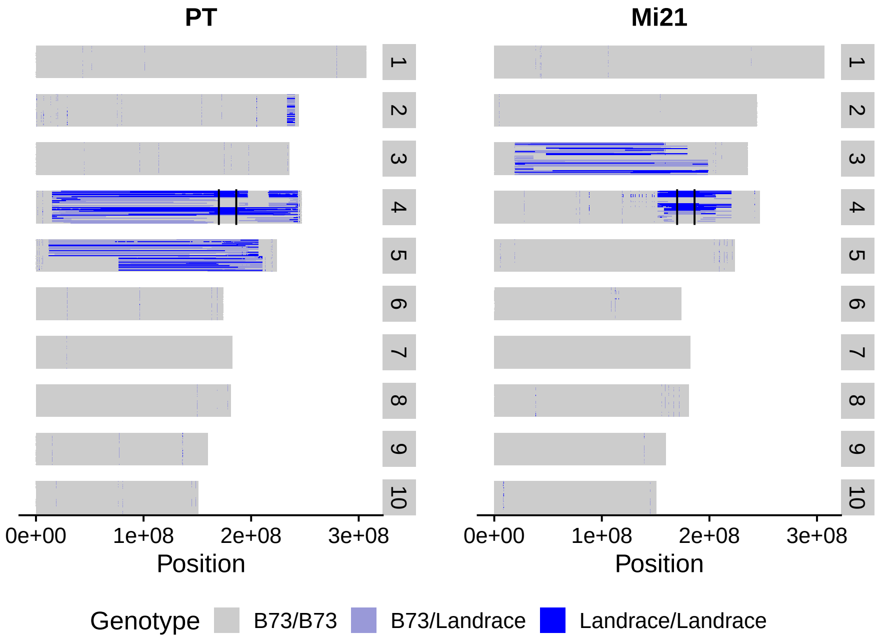


Figure 2: Genotyping results using RNA-seq data, and genotype correlation with *Inv4m*.

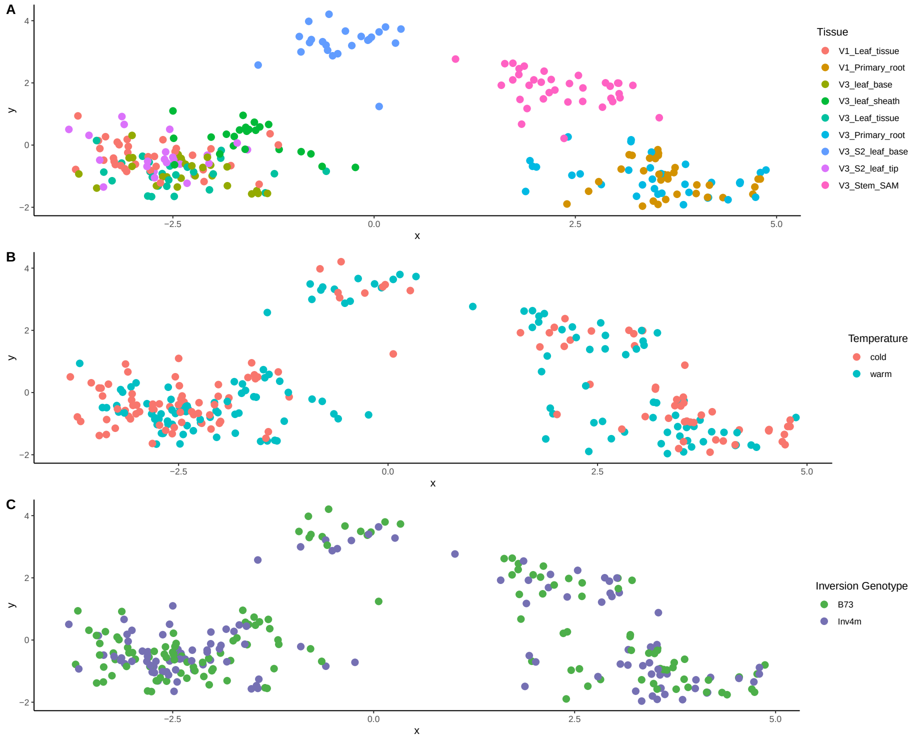


Figure 3: Multidimensional scaling plot of normalized and batch corrected gene expression (log2CPM) of the 500 highest expressed genes across our dataset using the *plotMDS* function. The plot represents a two-dimensional log2 fold change distance between each sample colored in three different ways to display how each factor effects the structure of the data. A) tissue B) *Inv4m* genotype and C) Temperature.

Table 1: Summary of linear mixed effects model of seedling emergence for each *Inv4m* donor and temperature treatment

| | | PT | | | Mi21 | | |
|---|---|---|---|---|---|---|---|
| Temperature | Predictors | Estimates | CI | p-value | Estimates | CI | p-value |
| Cold | Intercept | 9.55 | 7.49 – 11.62 | <0.001 | 8.94 | 7.97 – 9.91 | <0.001 |
| | Inv4m Genotype | -0.75 | -1.63 – 0.13 | 0.093 | -0.35 | -1.24 – 0.54 | 0.438 |
| | Observations | 40 | | | 51 | | |
| Warm | Intercept | 4.04 | 3.88 – 4.20 | <0.001 | 4.11 | 3.72 – 4.50 | <0.001 |
| | Inv4m Genotype | 0.07 | -0.09 – 0.23 | 0.396 | 0 | -0.27 – 0.27 | 0.989 |
| | Observations | 46 | | | 50 | | |

Table 2: Candidate genes within the *Inv4m* locus identified by their consistency of differential expression, correlation with *Inv4m-regulated* genes. Of the 89 genes identified, three were transcriptions factors, listed below.

| Location | List | Gene | Description |
|---|---|---|---|
| Inversion | Correlated Trans | Zm00001d051908 | OSIGBa0147B06.5 protein |
| Inversion | Correlated Trans | Zm00001d051998 | AP-1 complex subunit gamma-1 |
| Inversion | Correlated Trans | Zm00001d052075 | 60S ribosomal protein L35a-2 |
| Inversion | Correlated Trans | Zm00001d052079 | Lysine-ketoglutarate reductase |
| Inversion | Correlated Trans | Zm00001d052153 | YbaK/aminoacyl-tRNA synthetase-associated domain |
| Inversion | Correlated Trans | Zm00001d052259 | DCD (Development and Cell Death) domain protein |
| Inversion | Always DE | Zm00001d051872 | Aquaporin PIP1-5 |
| Inversion | Always DE | Zm00001d051882 | Ubiquitin carboxyl-terminal hydrolase |
| Inversion | Always DE | Zm00001d051987 | oxidoreductase zinc-binding dehydrogenase |
| Inversion | Always DE | Zm00001d052051 | Protein kinase superfamily protein |
| Inversion | Always DE | Zm00001d052136 | 1-aminocyclopropane-1-carboxylate oxidase20 |
| Inversion | Always DE | Zm00001d052210 | UDP-glycosyltransferase 85A7 |
| Inversion | Always DE | Zm00001d052242 | ATP synthase subunit delta chloroplastic |
| Inversion | Always DE | Zm00001d052245 | Exocyst complex component SEC6 |
| Inversion | Always DE | Zm00001d052269 | Putative auxin efflux carrier |
| Inversion | TF | Zm00001d051879 | NAC domain containing protein 71 |
| Inversion | TF | Zm00001d052180 | tunicate1 and MAPK cascade transcription positive |
| Inversion | TF | Zm00001d052229 | Ethylene-responsive transcription factor ERF056 |

Table 3: Candidate gene lists from 2 a priori gene pathway lists

| Location | List | Gene | Homolog | Function | Description |
|---|---|---|---|---|---|
| Trans | Flowering | Zm00001d028905 | PHYB | light transduction | Phytochrome |
| Trans | Flowering | Zm00001d045735 | COL1 | photoperiod | Constans1 |
| Trans | Flowering | Zm00001d044705 | TFL1 | integrator | |
| Trans | Flowering | Zm00001d031625 | SOC1 | integrator | mads77 - MADS-transcription factor 77 |
| Trans | Flowering | Zm00001d027425 | SOC1 | integrator | MADS-box transcription factor 56 |
| Trans | miRNA172 | Zm00001d002864 | | | |
| Trans | miRNA172 | Zm00001d026448 | | | Arabinose-proton symporter |
| Trans | miRNA172 | Zm00001d002075 | | | ereb212 - AP2-EREBP-transcription factor 212 |
| Trans | miRNA172 | Zm00001d039858 | | | ereb197 - AP2-EREBP-transcription factor 197) |
| Trans | miRNA172 | Zm00001d018409 | | | |
| Trans | miRNA172 | Zm00001d024635 | | | |
| Trans | miRNA172 | Zm00001d035512 | | | |
| Trans | miRNA172 | Zm00001d037377 | | | ereb81 - AP2-EREBP-transcription factor 81 |
| Trans | miRNA172 | Zm00001d002864 | | | |
